# The ‘Iceberg’ region of the Grapevine leafroll-associated virus 3 replicase polyprotein contains signals for mitochondrial targeting and outer membrane association

**DOI:** 10.1101/2025.09.05.674466

**Authors:** Catherine Fust, Caihong Li, Baozhong Meng

## Abstract

Positive-sense single-stranded (+ss) RNA viruses such as Grapevine leafroll-associated virus 3 (GLRaV3) replicate their genomes within membrane-bound viral replication complexes (VRCs). The biogenesis of such VRCs is driven by viral “replicase” polyproteins containing several replication-related domains. Past electron microscopy evidence suggests that GLRaV3 forms VRCs from the outer mitochondrial membrane of host grapevine plants. Here, we report the subcellular localization of the replicase polyprotein encoded by GLRaV3 *ORF1a* (PP1a) towards understanding its putative role in VRC formation. Through confocal laser scanning microscopy analysis of distinct EGFP-tagged PP1a domains, interdomain regions and truncations expressed in model plants, we report the molecular signals responsible for the targeting and association of PP1a to the mitochondria. This signal, located in the “Iceberg” region downstream from the methyltransferase-guanylyltransferase (M/GTase) domain, comprises an amphipathic α-helix and a downstream transmembrane domain (TMD). Mutagenesis studies suggest that the polar face of the amphipathic α-helix functions as the targeting signal, whereas the non-polar face, together with the TMD, act in membrane-anchoring. Microscopy observations are confirmed through mitochondrial isolation via gradient centrifugation and Western blotting. Structure prediction of the GLRaV3 M/GTase domain and its downstream TMD suggests a putative dodecameric oligomeric state. This dodecamer may gate the neck of GLRaV3 VRCs and contribute to their biogenesis, a hypothesis to be tested in follow-up studies.

**SIGNIFICANCE STATEMENT:** Positive-sense RNA viruses such as GLRaV3 form specialized membrane-bound viral replication complexes (VRCs) within host cells to sequester viral genome replication. Past evidence suggests that GLRaV3 targets the outer mitochondrial membrane (OMM) of grapevine host cells for VRC formation, however the viral protein responsible remained unknown. Here, we report a newly identified amphipathic α-helix and a downstream transmembrane domain in the replicase polyprotein encoded by GLRaV3 *ORF1a* that are crucial for OMM targeting and membrane association. Since OMM targeting mechanisms in plants are poorly understood, our findings not only shed light on how GLRaV3 assembles VRCs, but also provides insight into OMM targeting more broadly. This work lays the foundation towards elucidating the molecular mechanisms of GLRaV3 replication and host-pathogen interactions.

## INTRODUCTION

Grapevine leafroll-associated virus 3 (GLRaV3, species *Ampelovirus trivitis*) is the predominant agent of grapevine leafroll disease (GLRD), the most devastating viral disease of grapevines worldwide. GLRD reduces grapevine yield, negatively impacts berry development and quality, and decreases the productive lifespan of vineyards. In the absence of control measures such as early removal of symptomatic plants or insect vector management, GLRD can result in losses of up to $226,405 USD per vineyard hectare (Ricketts et al., 2015). Despite the severity of GLRaV3-induced GLRD and the worldwide distribution of the virus, little is known regarding its molecular biology and replication mechanisms (Burger et al., 2017; Fust et al., 2025).

GLRaV3 (genus *Ampelovirus*) belongs to the *Closteroviridae* family of viruses with 5’- capped, positive-sense, single-stranded (+ss) RNA genomes encapsidated within filamentous capsids ranging 650-2200 nm in length and 12 nm in diameter (Fuchs et al., 2020). Members of *Closteroviridae* possess the second largest +ssRNA genomes discovered to date ranging in size from 13-19 kb, only surpassed in size by some members of the *Coronaviridae*. *Closteroviridae* viruses share a conserved core of replication-related genes called the Replication Gene Block (RGB) comprised of two open reading frames (*ORFs*): *ORF1a* encoding PP1a, and *ORF1b* coding for PP1b (Koonin and Dolja, 1993). PP1a and PP1b replicase polyproteins are directly translated from viral genomic RNA upon entry and uncoating in a 5’ cap-dependent manner (Dolja et al.,1994; Agranovsky et al., 1994; Karasev 2000). Conserved domains in PP1a include a protease (PR) domain involved in polyprotein proteolytic processing (Peremyslov et al., 1998), a methyltransferase-guanylyltransferase (M/GTase) domain involved in RNA capping, and a RNA helicase (HEL) domain involved in RNA unwinding (Dolja et al., 2006). In contrast to other members of the *Closteroviridae* family, GLRaV3 and several other members of the *Ampelovirus* genus encode an Alkylation B (AlkB) domain within the translation product of *ORF1a* (Maree et al., 2008; Burger et al., 2017). This AlkB domain is believed to perform oxidative demethylation to protect viral +ssRNA genomes from methylation induced by host defense mechanisms (van den Born et al., 2008). The 5’ end of *ORF1b*, which encodes the RNA-dependent RNA polymerase (RdRP), overlaps with the 3’ end of *ORF1a*, and is translated into a PP1a/PP1b polyprotein following a +1 translational frameshift (Dolja et al.,1994; Agranovsky et al., 1994). The RdRP provides the polymerase activity required for the transcription of viral mRNAs and the replication of viral RNA genomes to generate progeny virions (Klaassen et al., 1996). All *ORFs* downstream from *ORF1b* are expressed through a nested set of subgenomic RNAs with 3’ co-linearity to the viral genome (Jarugula et al., 2010; Maree et al., 2010). These downstream *ORFs* include the second conserved gene block of *Closteroviridae*, the Quintuple Gene Block (QGB), spanning *ORFs 3-7* in GLRaV3 (Dolja et al., 2006). The QGB encodes proteins involved in virion assembly and cell-to-cell movement (Dolja et al.,1994; Dolja et al., 2006). Following the QGB is a unique gene block containing genes specific to each species of the family that vary in number and function.

Viruses with +ssRNA genomes such as GLRaV3 replicate their genomes within the cytoplasm of their host cell. These genome replication reactions are compartmentalized within virus-induced, membrane-bound, 50-300 nm diameter vesicles known as viral replication complexes (VRCs) (Unchwaniwala et al., 2021). VRCs are advantageous for viral replication because they shield double-stranded RNA genome replication intermediates from host defense mechanisms such as RNA silencing. Further, VRCs concentrate viral proteins and RNA to a confined space thereby enhancing efficiency in genome replication and transcription, and shields RNA synthesis from competing cellular processes such as translation. To form VRCs, select viral proteins target an intracellular membrane and induce membrane rearrangements in the form of single- or double-membraned vesicles (den Boon et al., 2010). Although the VRCs of varied +ssRNA viruses form from the membranes of various organelles such as the endoplasmic reticulum (ER), Golgi apparatus, or peroxisomes, individual +ssRNA virus species are selective in their membrane associations (Stapleford and Miller, 2010). Once formed, VRCs contain all viral proteins and host cell factors required for viral transcription and genome replication. These include viral replicase polyproteins (encoded by *ORF1a* and *ORF1b* for GLRaV3), viral genomic RNA, host ribonucleotides, among others. VRCs maintain a neck-like connection to host cell cytoplasm for the export of nascent viral mRNAs and genome copies, allowing for translation or genome packaging in host cell cytoplasm, respectively (Unchwaniwala et al., 2021). VRC necks are gated by “crown” complexes of viral replicase polyproteins with a central pore for RNA translocation.

Alphavirus supergroup viruses such as GLRaV3 possess an “Iceberg” region downstream from the M/GTase core domain containing membrane association sites conserved in structure, but not in sequence (Ahola and Karlin, 2015). This “Iceberg” region includes membrane-association sites involved in the anchoring of replicase crown complexes and the ensuing formation of VRCs, such as a conserved membrane-binding amphipathic α-helix (Ahola and Karlin, 2015; Unchwaniwala et al., 2021).

Electron microscopy (EM) studies of phloem specimens derived from GLRaV3-infected field-grown grapevine have revealed that GLRaV3 may form membrane-bound VRCs stemming from the outer mitochondrial membrane (OMM) of host cells (Kim et al., 1989; Faoro et al., 1991; Faoro and Carzaniga, 1995; Faoro, 1997). Given that the replicase polyproteins of +ssRNA viruses are the driving factor in VRC formation, this study aims to elucidate the subcellular localization and the predicted structure of the GLRaV3 PP1a replicase polyprotein towards understanding its putative role in VRC formation. Structural prediction of PP1a revealed an amphipathic α-helix and a downstream transmembrane domain (TMD) in the “Iceberg” region flanking the C-terminus of the M/GTase domain. This amphipathic α-helix and TMD were found to be necessary and sufficient to target mitochondria upon ectopic expression in the model plants. Mutagenesis revealed that the polar face of the amphipathic α-helix serves as the targeting signal, while the non-polar face along with the TMD serve as the membrane-associating “anchors”. To our knowledge, this research represents the first investigation of the subcellular localization of the GLRaV3 replicase polyprotein towards understanding its role in VRC formation.

## RESULTS

### In silico analysis of the replicase polyprotein encoded by GLRaV3 ORF1a reveals a putative amphipathic α-helix and TMD

The structure of the polyprotein encoded by GLRaV3 *ORF1a* (PP1a) was predicted using AlphaFold3, revealing that the known conserved domains (PR, M/GTase, AlkB, and HEL) are separated by extended intrinsically disordered regions (IDRs) (Fig. 1B), as evidenced by their low plDDT scores (Fig. 1C). The predicted structure of PP1a revealed a putative amphipathic α-helix within the Iceberg region of the M/GTase domain, spanning amino acids (AAs) 829-856 (Fig. 1A). This predicted amphipathic α-helix was further examined using HELIQUEST (Fig. 1D) (Gautier *et al*., 2008). The predicted amphipathic α-helix has a mean hydrophobicity of 0.65 and a hydrophobic moment of 0.41, suggesting that it is a strong candidate amphipathic α-helix.

**Figure 1:**
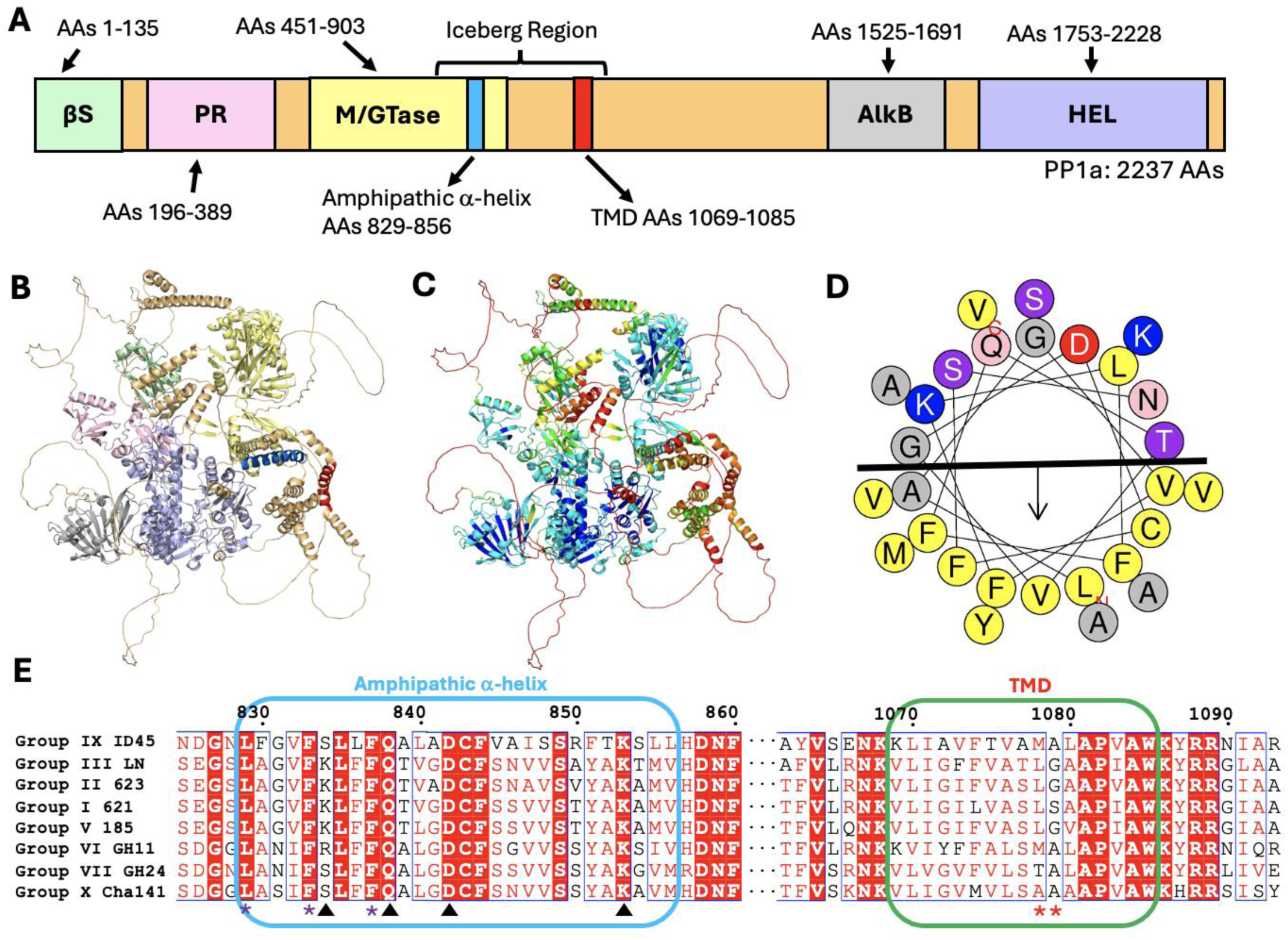
Bioinformatic analysis of GLRaV3 PP1a reveals an amphipathic α-helix and a downstream TMD in the Iceberg region. Schematic depiction of PP1a (A), predicted structure of PP1a colored by domain (B) or by plDDT score (C), helical wheel projection of the amphipathic α-helix (D), and conservation of the amphipathic α-helix and TMD among GLRaV3 isolates (E). The schematic depiction (A) and predicted structure (B) of GLRaV3 isolate 623 PP1a are colored by domain, with the βS domain depicted in green, the PR in pink, the M/GTase in yellow, the AlkB in grey, and the HEL in purple. IDRs are colored orange, while the amphipathic α-helix is depicted in blue, and the TMD in red. The predicted PP1a structure is colored by plDDT score (C) to discern prediction confidence. Residues with very low plDDT scores (<50) are coloured red, residues with low plDDT scores (50-60) are coloured orange, those with moderately low pIDDT scores (60-70) are coloured yellow, those with adequate plDDT scores (70-80) are coloured green, those with confident plDDT scores (80-90) are coloured cyan, and those with very confident plDDT scores (>90) are coloured blue. The helical wheel projection of the amphipathic α-helix (D) shows a clear hydrophobic face and hydrophilic face, separated by a black line. The arrow in (D) indicates the direction of the insertion into the cellular membrane. Absolutely conserved residues in the multiple sequence alignment (E) are shown as white text highlighted in red, and strongly conserved residues are displayed in red text and highlighted using blue boxes. Purple asterisks highlight amphipathic α-helix residues mutated in trM/GTaseΔLFF-trTMD, while black arrows highlight amphipathic α-helix residues mutated in trM/GTaseΔKQDK-trTMD, and red asterisks highlight TMD residues mutated in trM/GTase-ΔtrTMD.

The helical wheel projection of the amphipathic α-helix (Fig. 1D) demonstrates a clear hydrophobic face and hydrophilic face with a net positive charge, separated by a black line. The polar surface of the amphipathic α-helix would remain exposed to the cytoplasm, while its hydrophobic side could embed within one leaflet of the lipid bilayer, facilitating membrane association (Liu et al., 2009).

TOPCONS (Bernsel *et al.,* 2009) and TMHMM2.0 (Krogh *et al.,* 2001) identified a putative TMD downstream from the amphipathic α-helix of PP1a which spans AAs 1069-1085 (Fig. 1A). GLRaV3 isolates are divided into 9 phylogenetic groups due to sequence diversity (Diaz-Lara *et al.,* 2018). A multiple sequence alignment of the PP1a AA sequences of representative isolates from various variant groups (Fig. 1E) suggests that the TMD is conserved in all phylogroups and is flanked by conserved positively charged amino acids (AAs). Although the TMD sequence is not identical among isolates, its relative hydrophobicity as determined with the Kyte-Doolittle scale (Kyte and Doolittle, 1982) is under 2.0 and thus moderate in all isolates, suggesting conservation in function. TMDs of OMM-anchored proteins typically feature moderate to low hydrophobicity and a C-terminal extension of net positive charge (Lee *et al*., 2014; Sinning and McDowell, 2022), a property shared by the PP1a TMD.

Of interest, the predicted structure of PP1a revealed a putative domain that has not been reported in the literature. This new putative domain spanning AAs 1-135 forms a β-sandwich and will henceforth be named “βS” (Fig. 1A). The predicted βS domain lacks structural or sequence homologs in the protein database; we have therefore been unable to hypothesize its role in the replication and infection process of GLRaV3.

### Full-length GLRaV3 PP1a targets host mitochondria

Towards investigating the subcellular localization of GLRaV3 PP1a, full-length *ORF1a* was tagged to *EGFP* (Fig. 2A) and agroinfiltrated into leaves of *Nicotiana benthamiana* plants to mediate transient protein expression. PP1a-EGFP signal was visualized at 2 days post-infiltration (DPI) under confocal laser scanning microscopy (CLSM). When PP1a-EGFP was expressed alone (Fig. 2B), green punctate signal was readily observed in cells. When PP1a-EGFP was co-infiltrated along with a monomeric red fluorescent protein (mRFP)-tagged ER marker, mRFP-HDEL (Fig. 2C), punctate PP1a-EGFP signals were seen in close proximity to the ER network. The punctate PP1a-EGFP structures exhibited dynamic movement under CLSM (Movie S1), a property consistent with mitochondrial dynamics as they undergo frequent directional transport. Based upon the size, shape, and behaviour of the punctate structures, we postulated that they were most likely mitochondria. To pinpoint the GLRaV3 PP1a domains or structural features responsible for mitochondrial targeting, we next examined the subcellular localization of individual PP1a domains and interdomain regions.

**Figure 2:**
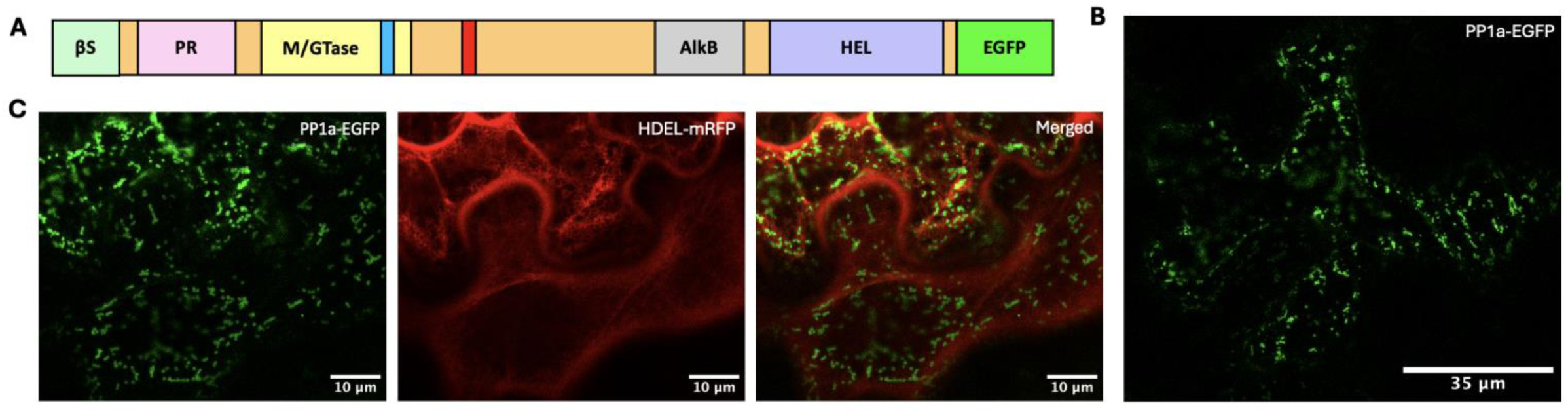
Mitochondrial localization of EGFP-tagged full-length GLRaV3 PP1a (PP1a-EGFP). Schematic depiction of the PP1a-EGFP construct (A), subcellular localization of PP1a-EGFP alone (B), and co-expression of PP1a-EGFP along with the HDEL-mRFP ER marker (C). All constructs were agroinfiltrated into model *N. benthamiana* and imaged at 2 days post-infiltration (DPI) via CLSM.

### The PP1a Iceberg region contains signals responsible for mitochondrial targeting and association

To identify PP1a regions or structural features that mediate mitochondrial targeting and association, eight constructs expressing EGFP-tagged PP1a domains and interdomain regions were generated (Fig. 3A), collectively spanning the entirety of PP1a. All constructs were transiently expressed in *N. benthamiana* plants and visualized at 2 DPI under CLSM. Six constructs lacked the bioinformatically-predicted amphipathic α-helix or TMD, namely βS-EGFP (AAs 1-135), PR-EGFP (AAs 196-389), βS-PR-EGFP (AAs 1-450), AlkB-EGFP (AAs 1524-1691), end of AlkB to the beginning of HEL (eAlkB-bHEL-EGFP, AAs 1692-1753), and HEL-EGFP (AAs 1754-2237). All six of these constructs localized to the cytosol (Fig. 3B) as evidenced by the diffuse signal displaying a pattern similar to free EGFP (Fig. 3C). The remaining constructs, M/GTase-EGFP (AAs 451-903) and TMD-EGFP (AAs 904-1523), contained the bioinformatically-predicted amphipathic α-helix or TMD respectively (Fig. 3A). While the TMD-EGFP construct encompassing all residues between the M/GTase and AlkB domains including the TMD and extensive IDRs localized to the ER (Fig. 3B), M/GTase-EGFP showed a localization pattern of particular interest. This latter construct displayed punctate signal reminiscent of mitochondria, in addition to diffuse cytosolic signal (Fig. 3B). We therefore concluded that M/GTase-EGFP containing the amphipathic α-helix was capable of targeting mitochondria albeit at lower levels.

**Figure 3:**
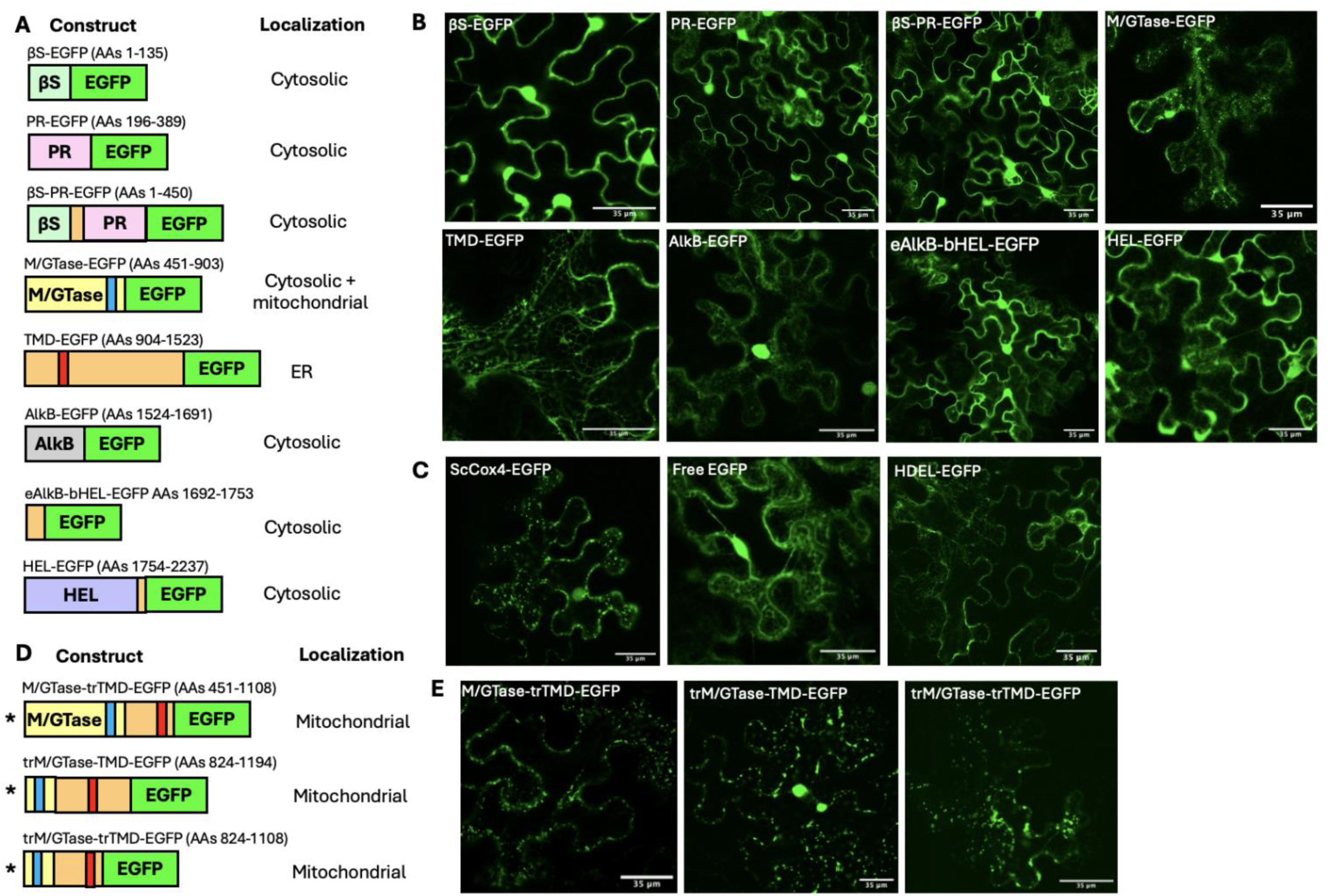
Subcellular localization of EGFP-tagged GLRaV3 PP1a domains, interdomain regions and truncations. (A) Schematic depiction of the EGFP-tagged GLRaV3 PP1a domain and interdomain region constructs generated towards investigating their subcellular localization. Eight C-terminal EGFP-tagged PP1a domain and interdomain region constructs were generated (A), with βS-EGFP including PP1a AAs 1-135, PR-EGFP AAs 196-389, βS-PR-EGFP AAs 1-450, M/GTase-EGFP AAs 451-903, TMD-EGFP AAs 904-1523, AlkB-EGFP AAs 1524-1691, eAlkB-bHEL-EGFP AAs 1692-1753, and HEL-EGFP AAs 1724-2237 (b = beginning of, e = end of). The subcellular localization of the proteins from constructs depicted in panel A is displayed in panel (B). Panel (C) depicts CLSM controls including the EGFP-tagged mitochondrial marker cytochrome C oxidase IV (ScCox4), free EGFP localizing to cytosol and diffusing into the nucleus, and the EGFP-tagged HDEL ER marker. Panel (D) displays schematic depictions of truncated GLRaV3 PP1a constructs featuring the bioinformatically-predicted amphipathic α-helix and TMD, namely M/GTase-trTMD-EGFP including PP1a AAs 451-1108, trM/GTase-TMD-EGFP AAs 824-1194, trM/GTase-trTMD-EGFP AAs 824-1108 (tr = truncated). (E) Subcellular localization of truncated constructs depicted in panel D. All constructs were agroinfiltrated into model *N. benthamiana* and imaged at 2 DPI via CLSM.

To investigate whether the mitochondrial localization of M/GTase-EGFP could be optimized, we extended the construct to include the downstream TMD of the Iceberg region. The resultant construct, M/GTase-truncated(tr)TMD-EGFP spanning AAs 451-1108 (Fig. 3D), displayed clear and abundant punctate localization (Fig. 3E) reminiscent of the signal of the EGFP-tagged mitochondrial marker cytochrome C oxidase IV (Fig. 3C). Towards fine-mapping the structural features responsible for efficient mitochondrial localization and association, two additional truncated constructs were generated. These truncated constructs, trM/GTase-TMD-EGFP (AAs 824-1194) and trM/GTase-trTMD-EGFP (AAs 824-1108) (Fig. 3D), still contained both the amphipathic α-helix and TMD. Both trM/GTase-TMD-EGFP and the further truncated trM/GTase-trTMD-EGFP displayed clear punctate signal similar in size, shape and behaviour to mitochondria (Fig. 3E).

To confirm organelle localization, co-infiltrations were performed along with the HDEL-mRFP ER marker. HDEL-mRFP signal co-localized with TMD-EGFP signal, confirming ER localization (Fig. S1A). HDEL-mRFP signal did not co-localize with trM/GTase-TMD-EGFP or trM/GTase-trTMD-EGFP, rather the EGFP signal from these two truncated constructs formed distinct punctate shapes which closely associated with the ER network, indicative of mitochondria (Fig. 4A and 4B). To further confirm mitochondrial localization of trM/GTase-TMD-EGFP and trM/GTase-trTMD-EGFP, these constructs were infiltrated into *Arabidopsis thaliana* plants transgenic for ATPaseΔ’PS-mRFP targeting mitochondria and imaged via CLSM at 3 DPI, the optimal time point for transient protein expression in *A. thaliana* according to Zhang *et al.,* (2020). Both trM/GTase-TMD-EGFP and trM/GTase-trTMD-EGFP co-localized with the mRFP-tagged mitochondria (Fig. 5A and 5B), while the localization of TMD-EGFP maintained an ER network-like appearance (Fig. S1B). Collectively, this data suggests that the amphipathic α-helix and TMD within GLRaV3 PP1a are minimally sufficient to mediate efficient mitochondrial targeting and association.

**Figure 4:**
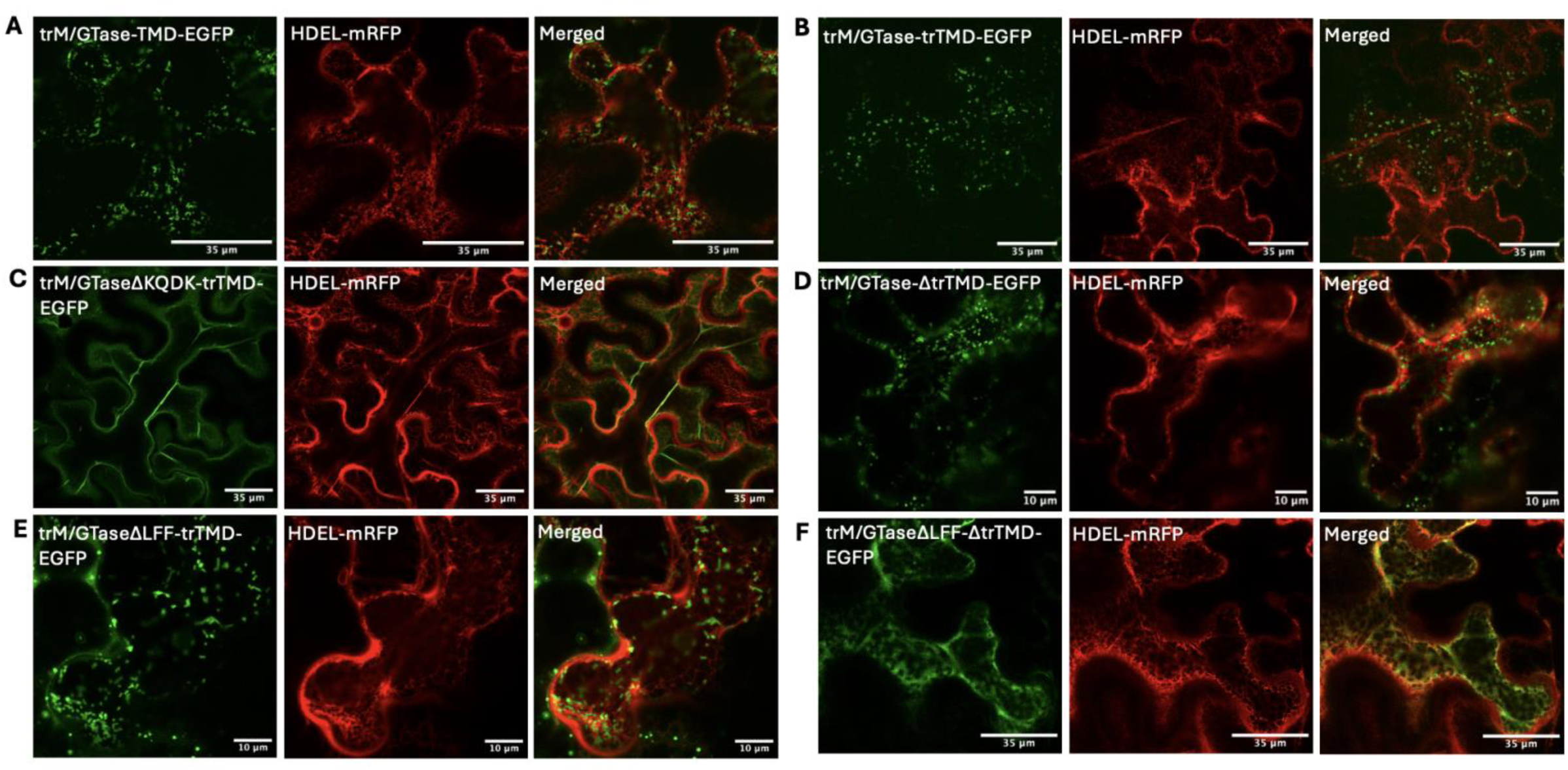
Co-expression of select EGFP-tagged GLRaV3 PP1a constructs with an HDEL-mRFP ER marker. Constructs were co-agroinfiltrated into leaves of *N. benthamiana* plants and imaged at 2 DPI via CLSM. The red channel displays the HDEL-mRFP signal, and the green channel displays the signal of the EGFP-tagged constructs. Merged images are provided to view co-localization, or lack thereof. Panel (A) displays signal from trM/GTase-TMD-EGFP, panel (B) trM/GTase-trTMD-EGFP, panel (C) trM/GTaseΔKQDK-trTMD, panel (D) trM/GTase-ΔtrTMD, panel (E) trM/GTaseΔLFF-trTMD, and panel (F) trM/GTaseΔLFF-ΔtrTMD.

**Figure 5:**
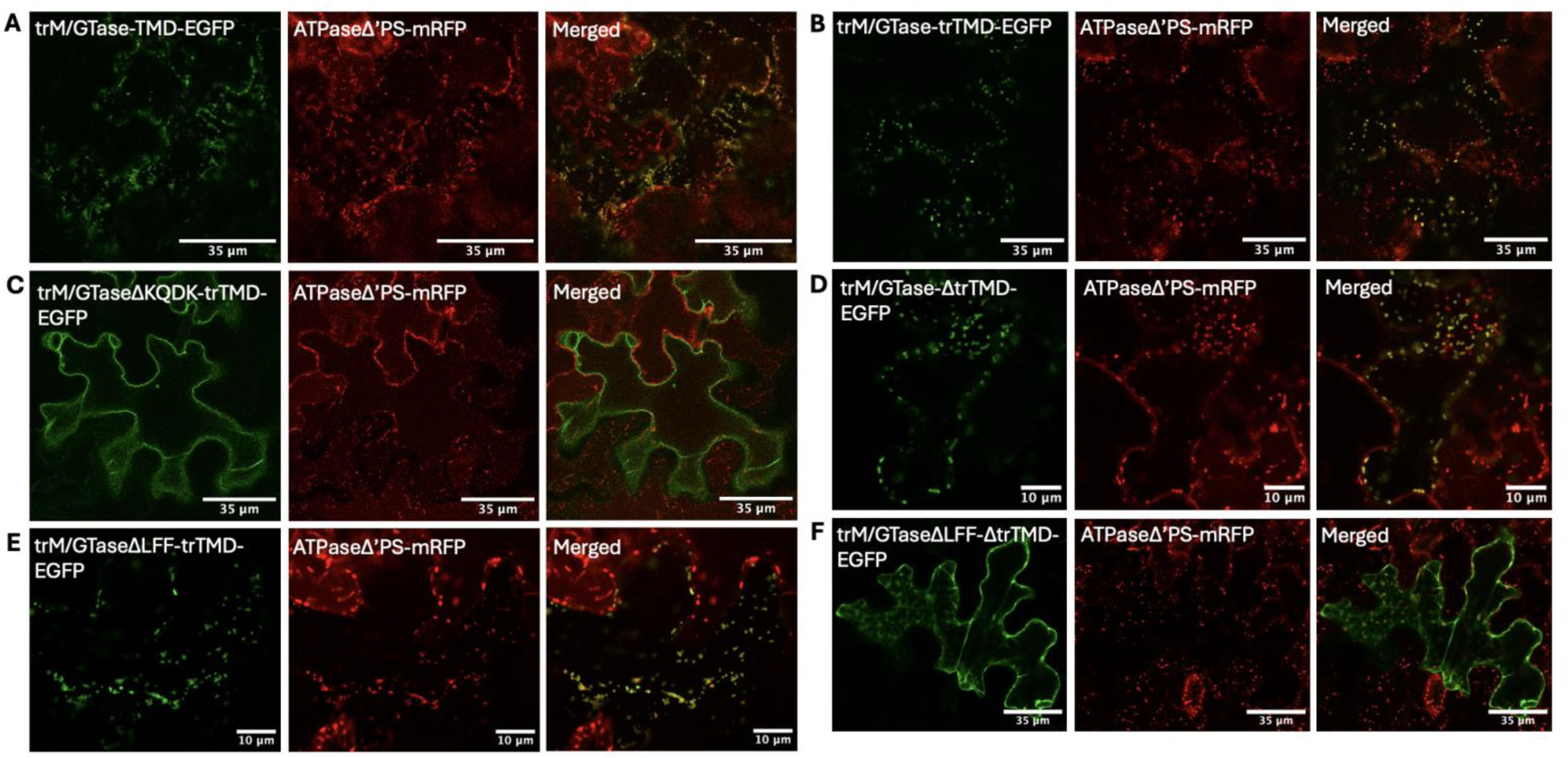
Expression of select EGFP-tagged GLRaV3 PP1a constructs in *A. thaliana* transgenic for ATPaseΔ’PS-mRFP. All constructs were agroinfiltrated into transgenic *A. thaliana* and imaged at 3 DPI via CLSM. The red channel displays the ATPaseΔ’PS-mRFP, whereas the green channel displays the signal of the EGFP-tagged constructs. Merged images are provided to view co-localization, or lack thereof. Panel (A) displays signal from trM/GTase-TMD-EGFP, panel (B) trM/GTase-trTMD-EGFP, panel (C) trM/GTaseΔKQDK-trTMD, panel (D) trM/GTase-ΔtrTMD, panel (E) trM/GTaseΔLFF-trTMD, and panel (F) trM/GTaseΔLFF-ΔtrTMD.

### The PP1a amphipathic α-helix and TMD are essential for mitochondrial targeting and association

Mutagenesis was performed to further dissect the role of the amphipathic α-helix and TMD of PP1a in mitochondrial targeting and membrane association. All mutations were performed in trM/GTase-trTMD-EGFP, the most truncated of the mitochondria-targeting constructs. To interrupt the polar face of the amphipathic α-helix, four conserved polar residues were mutated to non-polar residues, namely K834L, Q838L, D842L, and K853L, resulting in mutant construct trM/GTaseΔKQDK-trTMD-EGFP. This mutation successfully altered the localization pattern of the construct, leading to cytosolic signal (Fig. 4C, Fig. 5C). To interrupt the non-polar face of the amphipathic α-helix, three conserved non-polar residues were mutated to polar residues, namely L829E, F833E, and F837E, producing mutant construct trM/GTaseΔLFF-trTMD-EGFP. This mutant construct did not display an altered localization pattern, as punctate signal was still present when co-expressed with HDEL-mRFP (Fig. 4E), indicative of mitochondrial association. Further, the trM/GTaseΔLFF-trTMD-EGFP signal co-localized with the mRFP-tagged mitochondria of transgenic *A. thaliana* (Fig. 5E).

The next mutation aimed to interrupt the TMD of PP1a wherein two residues in the middle of the TMD, L1078 and G1079, were substituted to glutamate residues, producing mutant construct trM/GTase-ΔtrTMD-EGFP. Similarly to trM/GTaseΔLFF-trTMD-EGFP, the trM/GTase-ΔtrTMD-EGFP mutant produced punctate signal which lacked co-localization with HDEL-mRFP (Fig. 4D), but co-localized with ATPaseΔ’PS-mRFP (Fig. 5D). However, the combination of both the ΔLFF and ΔtrTMD mutations (trM/GTaseΔLFF-ΔtrTMD-EGFP) was sufficient to alter the localization pattern of the construct, leading to cytosolic signal (Fig. 4F, Fig. 5F).

### Biochemical analysis confirms the role of the PP1a amphipathic α-helix and TMD in mitochondrial targeting and association

To corroborate results from fluorescence microscopy and mutational analysis, we conducted biochemical analyses. Mitochondria were isolated from *N. benthamiana* plants expressing WT and mutant trM/GTase-trTMD-EGFP constructs at 2 DPI through fractionation with a Percoll gradient. Proteins extracted from the isolated mitochondria were subjected to Western blotting with an anti-GFP antibody. The Western blot (Fig. 6) corroborated the CLSM results, confirming mitochondrial localization of WT trM/GTase-trTMD-EGFP, trM/GTaseΔLFF-trTMD-EGFP, and trM/GTase-ΔtrTMD-EGFP as evidenced by a clear band of ∼57.5 kDa. These samples display a second band of ∼42 kDa (Fig. 6) which could represent a cleavage product. The blot lacks signal in mitochondrial proteins extracted from trM/GTaseΔKQDK-trTMD-EGFP and M/GTaseΔLFF-ΔtrTMD-EGFP, further suggesting that these mutations abolished mitochondrial targeting/association. Altogether, these results suggest that the polar face of the amphipathic α-helix acts as the OMM targeting signal, while the non-polar face of the amphipathic α-helix and TMD together act in membrane association.

**Figure 6:**
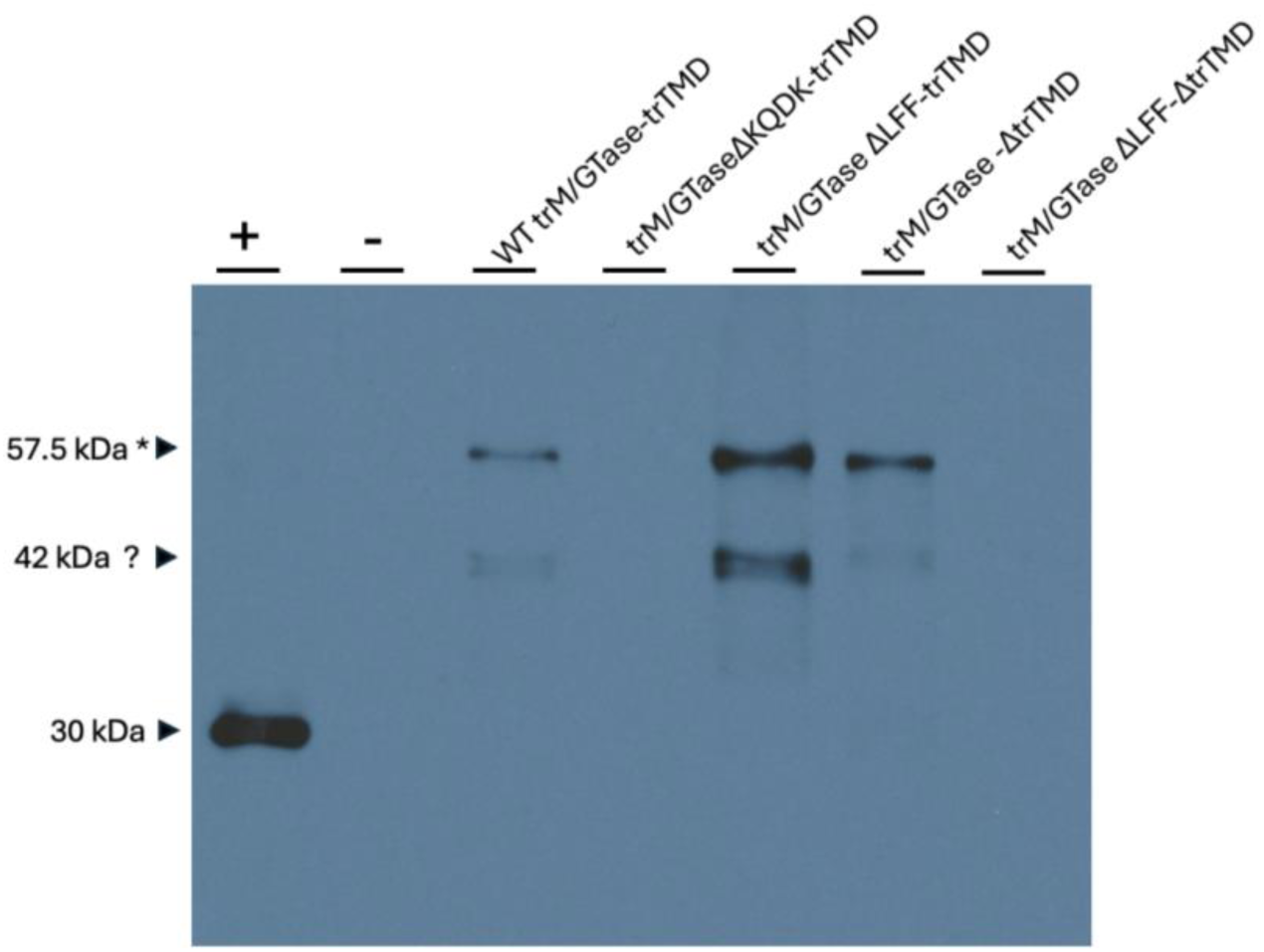
WT and mutant trM/GTase-trTMD mitochondrial localization confirmed by Western blot. Mitochondrial protein extracts from *N. benthamiana* plants expressing WT and mutant trM/GTase-trTMD constructs were analyzed by western blot with an anti-GFP antibody. The positive control (+) features a mitochondrial protein extract from a plant expressing an EGFP-tagged mitochondrial marker (N-terminal 29 AAs of *Saccharomyces cerevisiae* cytochrome oxidase IV), producing a band of ∼30 kDa. The negative control (-) features a mitochondrial protein extract from a plant expressing free EGFP, no signal is present. WT trM/GTase-trTMD, trM/GTaseΔLFF-trTMD, and trM/GTase-ΔtrTMD produce two bands, with the “*” band representing the expected size of ∼57.5 kDa. Mitochondrial protein extracts from plants expressing trM/GTaseΔKQDK-trTMD or trM/GTaseΔLFF-ΔtrTMD lack signal.

### The extended M/GTase domain of GLRaV3 may form a membrane-binding dodecamer

The structure of the GLRaV3 PP1a M/GTase domain plus its downstream Iceberg region (henceforth called extended M/GTase) was predicted with AlphaFold3. To our surprise, the resulting wedge-shaped structure of the extended M/GTase has strong structural homology (DALI Z-score of 24.3) to the experimentally determined structure of a dodecamer of the non-structural protein 1 (nsP1) complex of Chikungunya virus (CHIKV) (PDB 8APX) (Jones *et al.,* 2023). The extended M/GTase domain of PP1a was predicted as a trimer; four extended M/GTase trimers were superimposed onto 8APX to model a dodecamer (Fig. 7A-E). The resultant dodecamer of extended M/GTase domains of GLRaV3 PP1a places the amphipathic α-helix and the TMD in close proximity in the putative membrane-binding portion of the dodecamer (Fig. 7B and 7C). When colored by hydrophobicity (Fig. 7D), the amphipathic α-helix and TMD are shown to form an extended hydrophobic ring. Such a hydrophobic ring would be unstable in solution and as such would be buried within a membrane or protein interface, suggesting a potential role in membrane association. Examining surface electrostatics (Fig. 7E) reveals that the hydrophobic ring is flanked by an electropositive ring enriched in basic residues. This electropositive ring may interact favourably with phospholipid head groups, further hinting that the dodecamer may mediate membrane association. The central positively-charged channel of the dodecamer is enriched in basic residues and could coordinate RNA translocation (Fig. 7F) (Huang et al., 2024).

**Figure 7:**
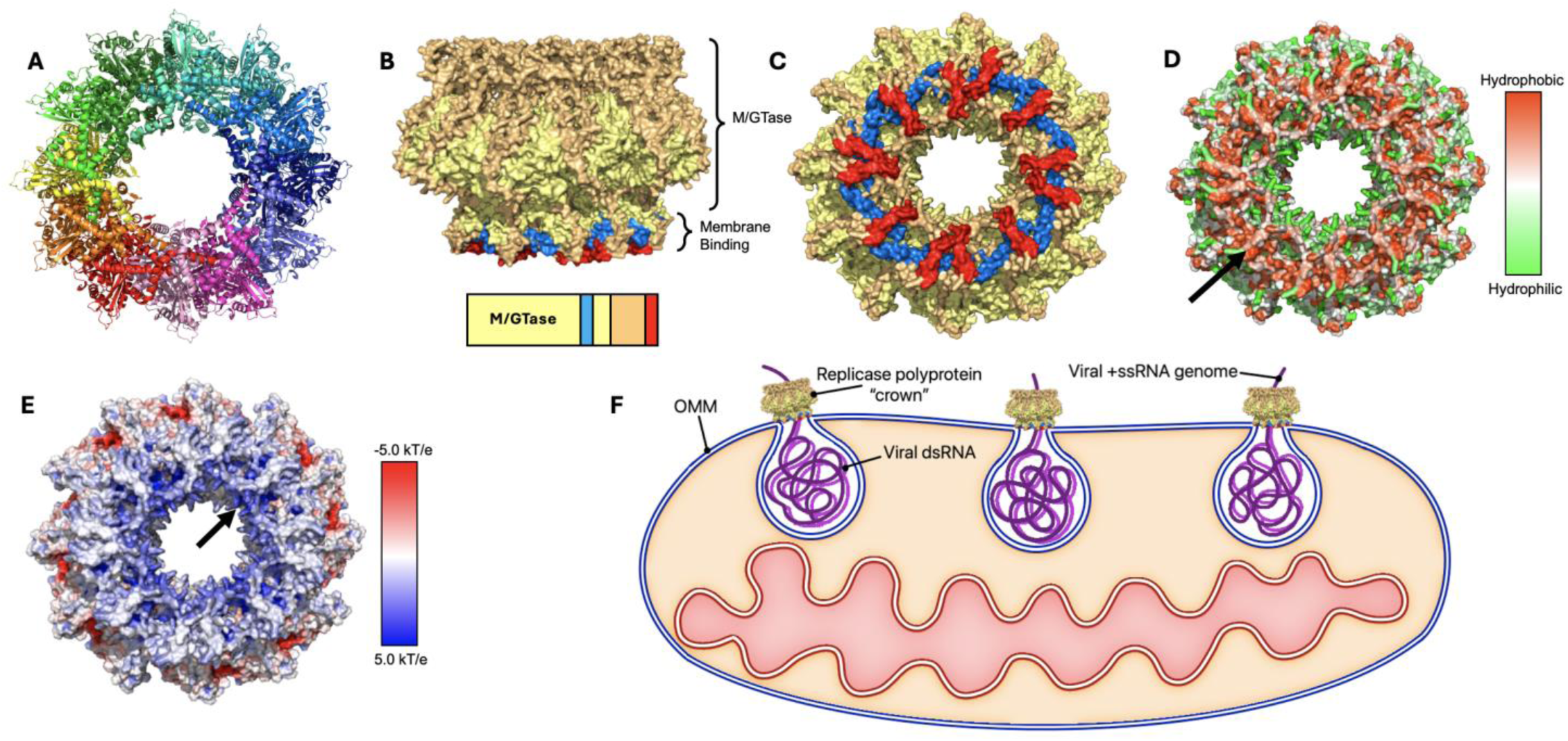
The extended M/GTase domain of GLRaV3 is predicted to form a OMM-binding dodecamer. Predicted dodecameric oligomeric state of the extended M/GTase domain of GLRaV3 PP1a (A-E) and schematic depiction of the dodecamer associating with the neck of OMM-derived VRCs (F). Panel (A) depicts the dodecamer of extended GLRaV3 M/GTase domains with each protomer displayed in a unique color, whereas (B) and (C) have the M/GTase domain in yellow, the amphipathic α-helix in blue, the intervening IDR in orange, and the TMD in red. The dodecamer is colored by residue hydrophobicity (D) with an arrow pointing to the hydrophobic (orange) ring and colored by electrostatics (E) with an arrow pointing to the electropositive (blue) ring.

## DISCUSSION

+ssRNA viruses are divided into three broad supergroups: the Alphavirus-like, the Picornavirus-like and the Flavivirus-like supergroups (Koonin 1991; Koonin et al., 2015). They take on distinct genome organization and expression strategies; however, the formation of membrane-bound VRCs is a phenomenon shared by +ssRNA viruses during their replication and infection process. Most well-studied +ssRNA viruses usurp the ER membrane of the host cell to form VRCs, while some viruses target organellar membranes such as the Golgi apparatus, peroxisomes, chloroplasts, or mitochondria (Liu et al., 2009; Gushchin et al., 2013). Preliminary EM evidence has suggested that GLRaV3 targets the OMM of grapevine host cells to form VRCs for viral transcription and genome replication (Kim et al., 1989; Faoro et al., 1991; Faoro and Carzaniga, 1995; Faoro, 1997). However, the mechanism of OMM targeting and association had yet to be identified.

Despite recent progress, protein targeting to the OMM remains poorly understood, and few +ssRNA viruses with OMM-associated replication have been studied. The notable exceptions include Carnation Italian ringspot virus (CIRV) and Flock House virus (FHV) which are distantly related to GLRaV3. While OMM targeting of CIRV relies upon its p36 replicase protein containing two moderately hydrophobic TMDs separated by an amphipathic α-helix with a positively charged polar face (Weber-Lotfi et al., 2002; Hwang et al., 2008), FHV relies upon an N-terminal TMD and an uncharacterized membrane association site located within the Iceberg region of its Protein A (Unchwaniwala et al., 2020). Given the evolutionary distance between GLRaV3 and both CIRV and FHV, and the limited study of viral OMM-targeting mechanisms, researching the Alphavirus-like supergroup member GLRaV3 may help reveal conserved OMM-targeting strategies shared by other +ssRNA viral proteins or even host proteins.

Our study demonstrates, for the first time, that full-length PP1a encoded by GLRaV3 *ORF1a* targets host mitochondria. Through investigating the subcellular localization of various EGFP-tagged PP1a domains, interdomain regions, and truncations, we have determined that the newly identified Iceberg region between AAs 824-1108 of PP1a, including the amphipathic α-helix and the downstream TMD, are minimally sufficient to mediate targeting to and association with the mitochondria. Specifically, mutagenesis revealed that the polar face of the amphipathic α-helix is required for targeting of mitochondria, as mutations to disrupt the polar face alone were sufficient to abolish mitochondrial localization. This is consistent with previous studies showing that positively charged polar faces of amphipathic α-helices contribute to OMM targeting (Hwang et al., 2008; Mason et al., 2014). Moreover, while disrupting either the non-polar face of the amphipathic α-helix or the TMD individually was insufficient to block mitochondrial association, the combination of both mutations effectively abolished this interaction. This indicates that, at least within the trM/GTase-trTMD construct, the non-polar face of the amphipathic α-helix and the TMD exhibit functional redundancy in OMM anchoring. While these elements appear to serve overlapping functions in this small, truncated construct, it is important to note that the precise cleavage pattern of PP1a remains undefined. The cleavage product containing the Iceberg region is likely substantially larger and may adopt a dodecamer as its oligomeric state. It is likely that both the non-polar face of the amphipathic α-helix and the TMD are necessary to anchor the larger multimeric complex to the OMM. Of course, this is an idea that warrants further investigation. Findings from this research reveal that GLRaV3 shares certain aspects with CIRV and FHV in relation to localization to the OMM despite that they are very different from one another in genome structure, expression strategy and hosts. For instance, all three viruses require 1-2 TMDs for anchoring in the OMM. However, while FHV Protein A employs signal-anchor mechanism with its N-terminal TMD (Miller and Ahlquist, 2002) and CIRV p36 employs a signal loop-anchor mechanism (Hwang et al., 2008), the extended M/GTase domain of GLRaV3 would likely be tail-anchored upon cleavage by the viral leader protease. This represents a potentially new and distinct mechanism for protein targeting to the OMM.

The dodecameric crown model of the extended GLRaV3 M/GTase domain assumes that cleavage occurs, at least, between the PR and M/GTase domains, and between the M/GTase and AlkB domains. Although the PR domain of PP1a is predicted to autocatalytically cleave between the PR and M/GTase domains at Gly371/Gly372 (Ling *et al.,* 2004; Mostert *et al*., 2023), experimental evidence for this and other possible cleavage sites remains lacking and warrants future investigation. Nonetheless, the strong structural homology between CHIKV 8APX and the extended GLRaV3 M/GTase domain is compelling. The atomic structure of dodecameric, membrane-binding M/GTase crowns involved in VRC formation have been recently resolved in vertebrate-infecting CHIKV (8APX) (Jones et al., 2023) and invertebrate-infecting FHV (Unchwaniwala et al., 2020), but no analogous structure has yet been reported for any plant virus. The dodecameric crowns formed by CHIKV M/GTase were shown to recruit HEL and RdRP domains to drive viral genome replication (Tan et al., 2022). The RdRP activity and active viral RNA synthesis of the crown complex is thought to contribute to vesicle formation, as the growing dsRNA gradually fills the vesicle interior (Unchwaniwala et al., 2021).

Investigating protein-protein interactions among the various domains from the cleavage of polyproteins encoded by GLRaV3 may therefore provide valuable insights into the biogenesis of VRCs. Further work may include employing cryo-EM on mitochondria isolated from GLRaV3-infected plants to visualize VRCs to further support their organellar origin, and investigating interactions among different domains of the replicase polyproteins. Additionally, since host proteins involved in lipid synthesis, membrane modification, and membrane curvature are often recruited to VRCs (Aktepe et al., 2017; Ketter et al., 2019; Zhang et al., 2019), identifying host proteins co-opted by GLRaV3 would provide valuable insights into VRC biogenesis in particular, and pathogenesis of GLRaV3 infection in the grapevine host in general. Lastly, because mitochondria play crucial roles in plant immunity, including in programmed cell death (Wang et al., 2022), the potential impact of GLRaV3 on host immune responses warrants investigation. Viral disruption of mitochondrial function has been shown to modulate immune mechanisms upon infection with flaviviruses (Losarwar et al., 2025) and coronaviruses (Gatti et al., 2020), as well as with plant viruses such as melon necrotic spot virus (Navarro et al., 2021) and rice ragged stunt virus (Zheng et al., 2025).

In closing, our study demonstrates that the Iceberg region of GLRaV3 PP1a harbors critical signals for mitochondrial targeting and OMM association. Specifically, the hydrophilic face of the amphipathic α-helix governs mitochondrial localization, while its hydrophobic face, together with the downstream TMD, ensures anchoring to the OMM. Structural modeling further reveals that GLRaV3 M/GTase domains and Iceberg regions may self-assemble into a dodecameric crown-like structure, strikingly reminiscent of the recent cryo-EM-resolved architectures in FHV (an invertebrate virus) and CHIKV (a human-infecting vertebrate virus).

This structural conservation across diverse RNA viruses underscores a potentially ancient and universal mechanism in the VRC formation across vast evolutionary distances. These findings provide a solid foundation for future investigations into the molecular, cellular, and pathological mechanisms underpinning GLRaV3 replication and pathogenesis, with broader implications for understanding RNA virus biology and developing targeted antiviral strategies.

## MATERIALS AND METHODS

### In silico analysis of GLRaV3 PP1a

The AA sequence of GLRaV3 PP1a (isolate 623) was retrieved from NCBI (ADI49425.1). The entire 2237-AA sequence of GLRaV3 PP1a was predicted with AlphaFold3 (Abramson *et al.,* 2024). The domains were colored in PyMOL. When colored by residue hydrophobicity, an amphipathic α-helix within PP1a became evident. This amphipathic α-helix was further examined in HELIQUEST (Gautier *et al*., 2008). TOPCONS (Bernsel *et al.,* 2009) and TMHMM2.0 (Krogh *et al.,* 2001) were employed to locate putative transmembrane domains (TMDs). A multiple sequence alignment of the PP1a TMD AA sequences of various GLRaV3 isolates representing each phylogroups was performed with CLUSTALW (Kyoto University Bioinformatics Center) and presented using ESPript 3.0 (Robert and Gouet, 2014).

### Plasmid construction of EGFP-tagged ORF1a segments and site-directed mutagenesis

Full-length GLRaV3 *ORF1a* or *ORF1a* domains and inter-domain regions were initially cloned into pRTL2m4-*eGFP* vectors to enable in-frame tagging with *EGFP*. The MCS is preceded by duplicated enhancer, a *35S* promoter from Cauliflower mosaic virus (CaMV) to promote constitutive transcription in all plant tissues, and a translational leader sequence from Tobacco etch virus (TEV). The vector contains a CaMV polyadenylation signal and a nopaline synthase terminator sequence downstream from the MCS.

Quick-change site-directed mutagenesis (Liu and Naismith, 2008) was employed to introduce a silent mutation to PP1a of strain 623 towards removing an internal *Nco*I restriction site (Supplementary Table 1). GLRaV3 *ORF1a* domains and inter-domain regions were PCR-amplified from the modified GLRaV3 strain 623 genome with gene-specific primers (Supplementary Table 1). PCR was conducted with high-fidelity PaCeR polymerase (GeneBioSystems) according to manufacturer’s instructions. A *Nco*I restriction site was included in the forward primer, and a *Kpn*I restriction site in the reverse primer. Amplified fragments were gel-purified using GenepHlow™ Gel/PCR Kit (Geneaid). pRTL2m4-*EGFP* and the amplified fragments were digested *Nco*I and *Kpn*I FastDigest restriction enzymes (Thermo Fisher Scientific) according to manufacturer’s instructions. *ORF1a* domains and inter-domain regions were ligated into pRTL2m4-*EGFP* using T4 DNA ligase (New England Biolabs) according to the manufacturer’s instructions, and subsequently transformed into heat-shock treated chemically competent *Escherichia coli* JM109. Positive transformant colonies were selected following 16-hour 37 °C growth on LB-Ampicillin plate (50 µg/mL) and confirmed by colony PCR. Plasmids were isolated from overnight cultures with a Presto™ Mini Plasmid Kit (Geneaid) according to manufacturer’s instructions.

Constructs were next subcloned into binary pCAMBIA-0380 vectors. pCAMBIA-0380 and the pRTL2m4-*EGFP* constructs were digested with FastDigest *Hind*III and *Bcu*I (Thermo Fisher Scientific) restriction enzymes, gel extracted and ligated as previously described. Positive colonies were selected on LB-Kanamycin plates (50 µg/mL) and isolated as previously described. Cloning success was further confirmed through Sanger sequencing at the University of Guelph Advanced Analysis Centre (AAC). Once validated, plasmids were transformed into *Agrobacterium tumefaciens* strain EHA105.

Quick-change site-directed mutagenesis (Liu and Naismith, 2008) was employed to introduce ΔKQDK, ΔLFF and/or ΔtrTMD mutations in pCAMBIA vectors containing trM/GTase-trTMD-EGFP (Supplementary Table 1). Mutagenesis success was further confirmed through Sanger sequencing. Once validated, plasmids were transformed into *A. tumefaciens* strain EHA105.

### Plant material and growth conditions

All plants were grown under controlled conditions in Conviron growth chambers at the University of Guelph’s Phytotron facility. The conditions included 14 hours of 120 μmole intensity light at 24°C, to 10 hours of dark at 21°C with 60% humidity.

Wildtype *Nicotiana benthamiana* model plants seeds were sown on moistened PRO-MIX^®^ BX (Premier Tech Horticulture) soil with a humidity cover until germination. 2 weeks post-sowing, *N. benthamiana* plantlets were transferred to 3 ½” by 3 ½” pots containing PRO-MIX^®^ BX soil moistened with fertilized water, and a humidity cover was placed for an additional week. 3 weeks post-sowing, humidity covers were gradually removed from the plantlets over the course of 2 days to allow for gradual exposure to lower humidity levels. Once humidity covers were removed, the plants were watered every 3 days. 6-week-old *N. benthamiana* plants were employed for transient expression studies.

Transgenic *Arabidopsis thaliana* expressing an mRFP-tagged ATPaseΔ’PS targeting the mitochondria (kindly provided by Dr. S. Arimura from the University of Tokyo, Japan) were grown as described above for *N. benthamiana*. 5-week-old transgenic *A. thaliana* plants were employed for transient expression studies.

### Agroinfiltration to mediate transient protein expression

For agroinfiltration into *N. benthamiana* model plants, glycerol stocks of *Agrobacterium tumefaciens* strain EHA105 containing plasmids-of-interest were streaked onto LB plates supplemented with Rifampicin (10 µg/mL) and Kanamycin (50 µg/mL) and incubated at 30°C for 48 hours to grow colonies. *A. tumefaciens* colonies were picked and grown in 5 ml liquid LB cultures containing Rifampicin (10 µg/mL) and Kanamycin (50 µg/mL) for 18 hours at 30°C with 150 RPM agitation. 3 ml of overnight culture was sub-cultured in 35 ml of fresh liquid LB with Rifampicin and Kanamycin and grown for approximately 3.5 hours at 30°C to an optical density 600 (OD_600_) of 1.0. Cultures were then pelleted (5000 RPM, 6 minutes), and supernatants were discarded. Pellets were resuspended in 10 ml of infiltration buffer (10 mM MES, 10mM MgSO_4_), pelleted once again (5000 RPM, 6 minutes), and the wash supernatant was discarded.

Pellets were next resuspended in infiltration buffer to an OD_600_ of 0.2 and supplemented with 150 µM acetosyringone for induction. Cells sat at room temperature in the induction buffer for 20 hours prior to agroinfiltration, following the procedure outlined by Lindbo *et al*. (2007).

Briefly, cells were infiltrated into the abaxial surface of *N. benthamiana* leaves using a 1 ml needleless syringe. Bacterial growth, preparation and agroinfiltration into transgenic *A. thaliana* model plants was carried out as described in Zhang *et al.,* (2020), without modification.

### Confocal laser scanning microscopy

Confocal laser scanning microscopy (CLSM) was conducted with an upright Leica DM 6000B microscope connected to a Leica TCS SP5 system with a 488 nm Argon laser and a 543 nm He-Ne laser. Emission spectra of 503-524 nm and of 566-643 nm were employed to image EGFP-tagged proteins and mRFP-tagged proteins, respectively. CLSM employed *N. benthamiana* plants expressing constructs-of-interest at 2 DPI (Lindbo *et al*. 2007), and *A. thaliana* plants expressing constructs-of-interest at 3 DPI (Zhang *et al.,* 2020). Microscopy sample preparation involved excising 1.5 cm^2^ samples of a leaf-of-interest with a scalpel and mounting the sample onto a microscope slide under a cover slip with water as a mounting medium. Images were processed in Fiji.

### Isolation of mitochondria from N. benthamiana

Mitochondria were isolated from *N. benthamiana* cells expressing constructs-of-interest with a discontinuous Percoll centrifugation media (Cytiva) gradient as described by Gualberto *et al*. (1995) with some modifications. 20 g of *N. benthamiana* leaves 2 days post-infiltration with constructs-of-interest were homogenized with 50 ml of extraction buffer (0.4 M sucrose, 50 mM Tris-HCl (pH 7.5), 3 mM EDTA (pH 8.0), 0.2 mM EGTA, 0.1% BSA, and 4 mM β-mercaptoethanol) in a waring blender. All subsequent steps were carried out on ice. The homogenates were filtered through 2 layers of cheesecloth into sterile 50 ml centrifuge tubes and centrifuged at 3,000 g at 4°C for 10 mins. Supernatants were placed into sterile 40 ml PPCO centrifuge tubes (Nalgene™) and centrifuged at 10,000 g at 4°C for 20 mins with a JA-25.50 rotor. Supernatants were discarded, and pellets were resuspended in 12 ml of extraction buffer in a dounce homogenizer. Resuspended pellets were transferred back into the 40 ml PPCO centrifuge tubes and centrifuged at 1,000 g at 4°C for 5 mins. Supernatants were placed into new 40 ml PPCO centrifuge tubes and centrifuged at 10,000 g at 4°C for 15 mins. Supernatants were discarded, and pellets were resuspended in 1 ml of extraction buffer. The resuspension was layered on top of a 10 ml discontinuous Percoll gradient (2 ml 45% Percoll, 4 ml 21% Percoll, 4 ml 13% Percoll) in 13.2 ml Ultra-Clear™ tubes (Beckman Coulter, 344059) and centrifuged at 13,000 g at 4°C for 30 mins with a SW 41 rotor. Mitochondria sediment in a translucent band in the bottom 1 ml of the tube, these bands were collected with a Pasteur pipette and placed into clean 40 ml PPCO centrifuge tubes. Isolated mitochondria were washed in 12 ml of extraction buffer to remove remaining Percoll and centrifuged at 10,000 g at 4°C for 15 mins. The supernatant was carefully removed, and mitochondrial pellets were washed three additional times in extraction buffer without BSA and β-mercaptoethanol. Following washes, mitochondria were pelleted and resuspended in 50 µl of extraction buffer without BSA and β-mercaptoethanol.

### Western blotting

16.67 µl of each mitochondrial extract-of-interest was mixed with 2.78 µl of 6X SDS loading buffer and boiled for 10 minutes to denature mitochondrial proteins. SDS-PAGE, wet transfer to PVDF and membrane blocking were carried out as described by Meng et al., (2003). All subsequent steps were carried out at room temperature unless otherwise specified. Blocked membranes were incubated with 1:4000 primary anti-GFP N-terminal rabbit antibody (Sigma-Aldrich) in 1X PBS-T containing 1% skim milk overnight at 4°C with 60 RPM agitation. Membranes were washed with 25 ml of 1X PBS-T four times at 5 minutes each with 60 RPM agitation to remove unbound primary antibody. Membranes were incubated with 1:8000 secondary ECL™ anti-rabbit IgG, Horseradish peroxidase-linked whole antibody from donkey (GE Healthcare) in 1X PBS-T containing 1% skim milk for 2 hours with 60 RPM agitation. Membranes were washed as described previously. Washed and drained membranes were incubated with 2 ml of Pierce™ ECL Western Blotting Substrate (Thermo Fisher Scientific) for 5 minutes, and protein bands were exposed to X-ray films for 2 minutes to capture signal.

### Determining the putative oligomeric state of the extended M/GTase domain of GLRaV3 PP1a

The AA sequence of PP1a from GLRaV3 isolate 623 was downloaded from NCBI (ADI49425.1). Residues 451-1105 corresponding to the M/GTase domain and downstream TMD were submitted to AlphaFold3 (Abramson *et al.,* 2024) for structural prediction of a trimer. The rank 0 trimeric structure was employed for downstream analysis. A DALI protein structure comparison search was performed against the complete Protein Data Bank (PDB) database to identify homologous structures, with the closest hit being a dodecamer of the Chikungunya virus (CHIKV) non-structural protein 1 (nsP1) complex (PDB 8APX) (Jones *et al.,* 2023). To model a dodecamer of the extended M/GTase domain of GLRaV3, four trimeric extended M/GTase structures were superimposed onto 8APX using DALI and visualized in PyMOL.

## ACKNOWLEDGMENTS

The authors wish to acknowledge Dr. M. Strüder-Kypke for her guidance with CLSM at the Molecular and Cellular Imaging Facility at the University of Guelph. We thank Dr. J. Mathur and Dr. R. Mullen for fluorescent probes, Dr. M. Kimber for support with bioinformatic analyses, Dr. S. Arimura for supplying transgenic *A. thaliana*, and Dr. L. Rubino for the mitochondrial isolation protocol. Finally, we thank G. Jackson for her assistance with the cloning of a construct.

---

**Supplementary Figure 1:**
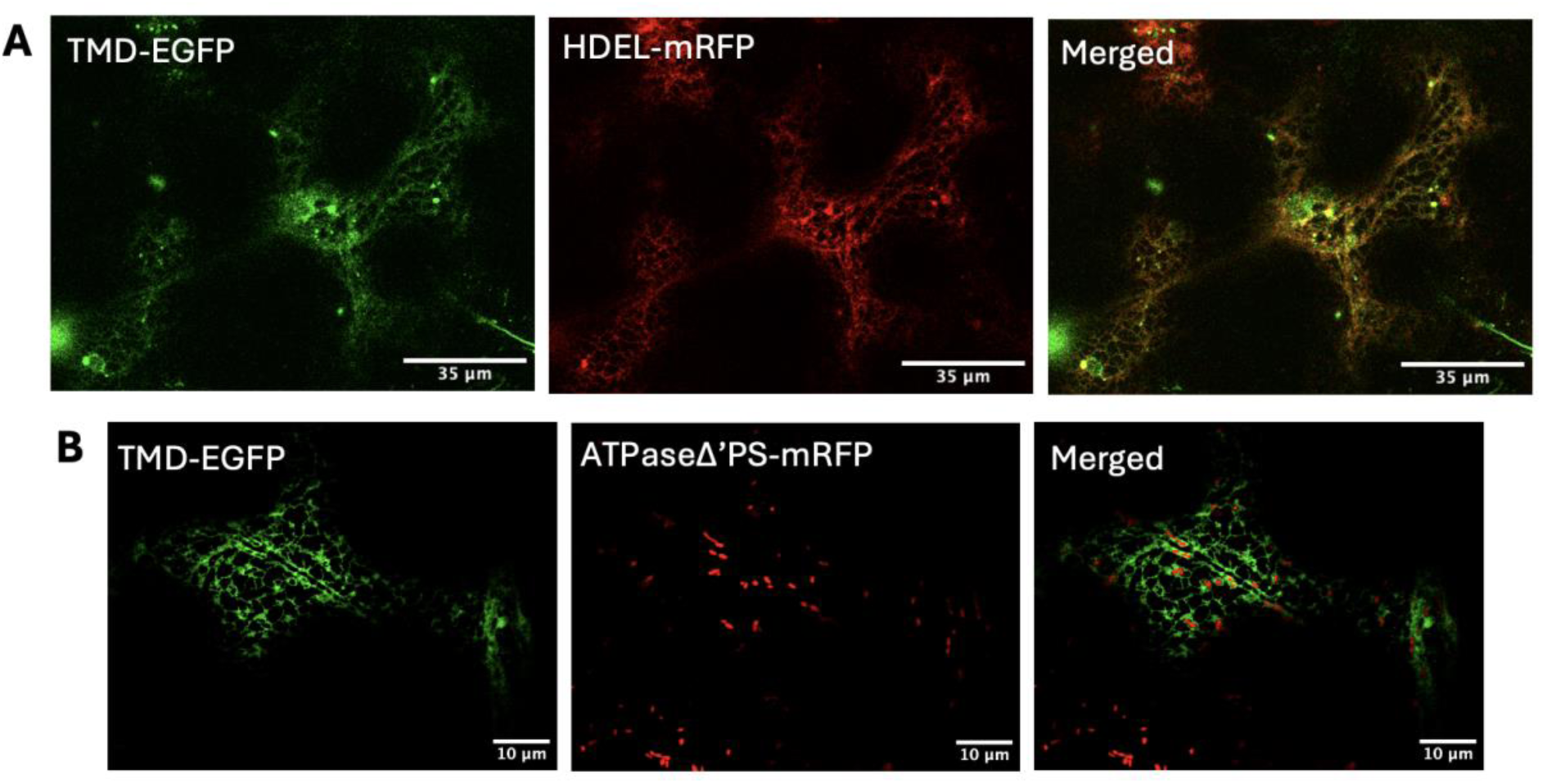
ER localization of TMD-EGFP. TMD-EGFP co-localizes with HDEL-mRFP (A) when co-infiltrated in *N. benthamiana*. When expressed in *A. thaliana* transgenic for ATPaseΔ’PS-mRFP, the mRFP-tagged mitochondrial signal is closely associated with the TMD-EGFP signal but lacks co-localization.

**Supplementary Table 1:**
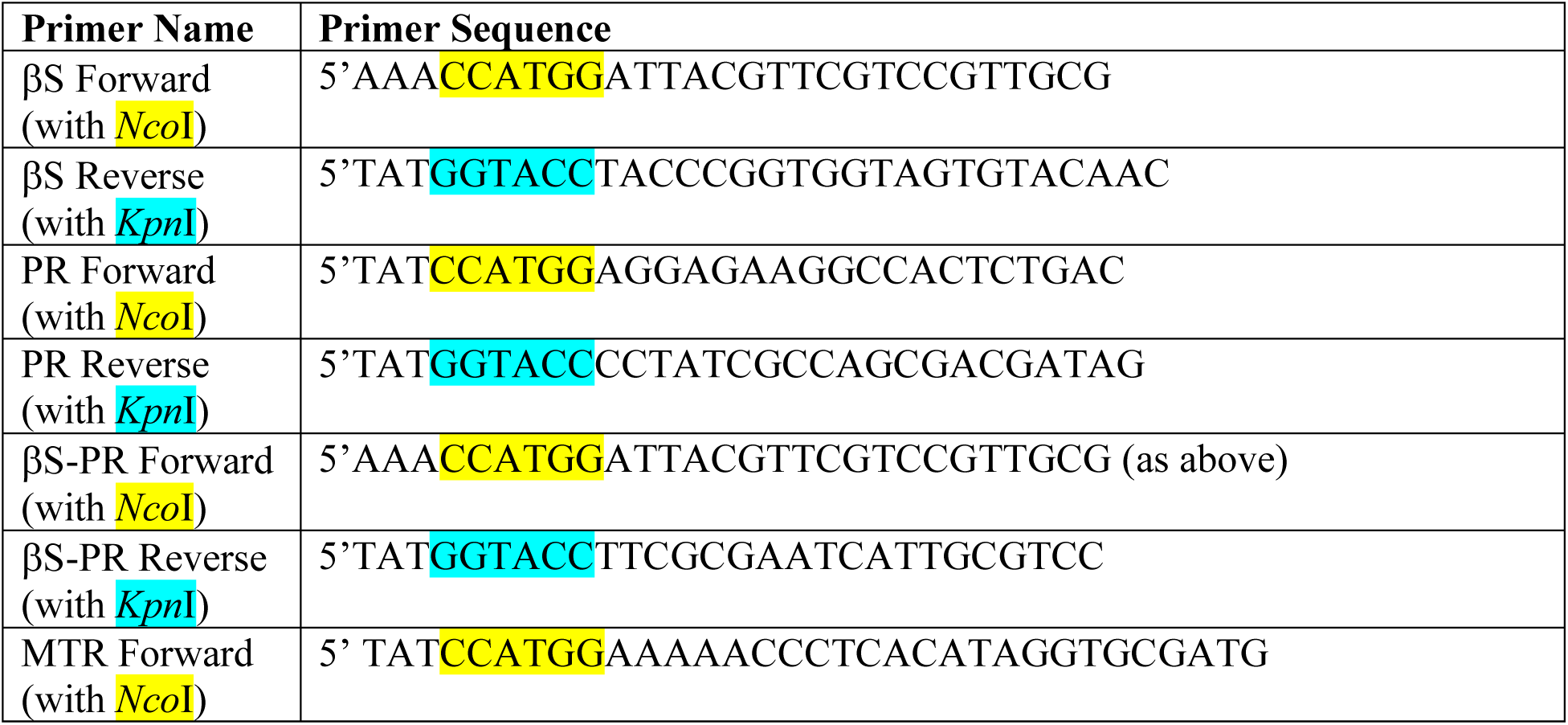
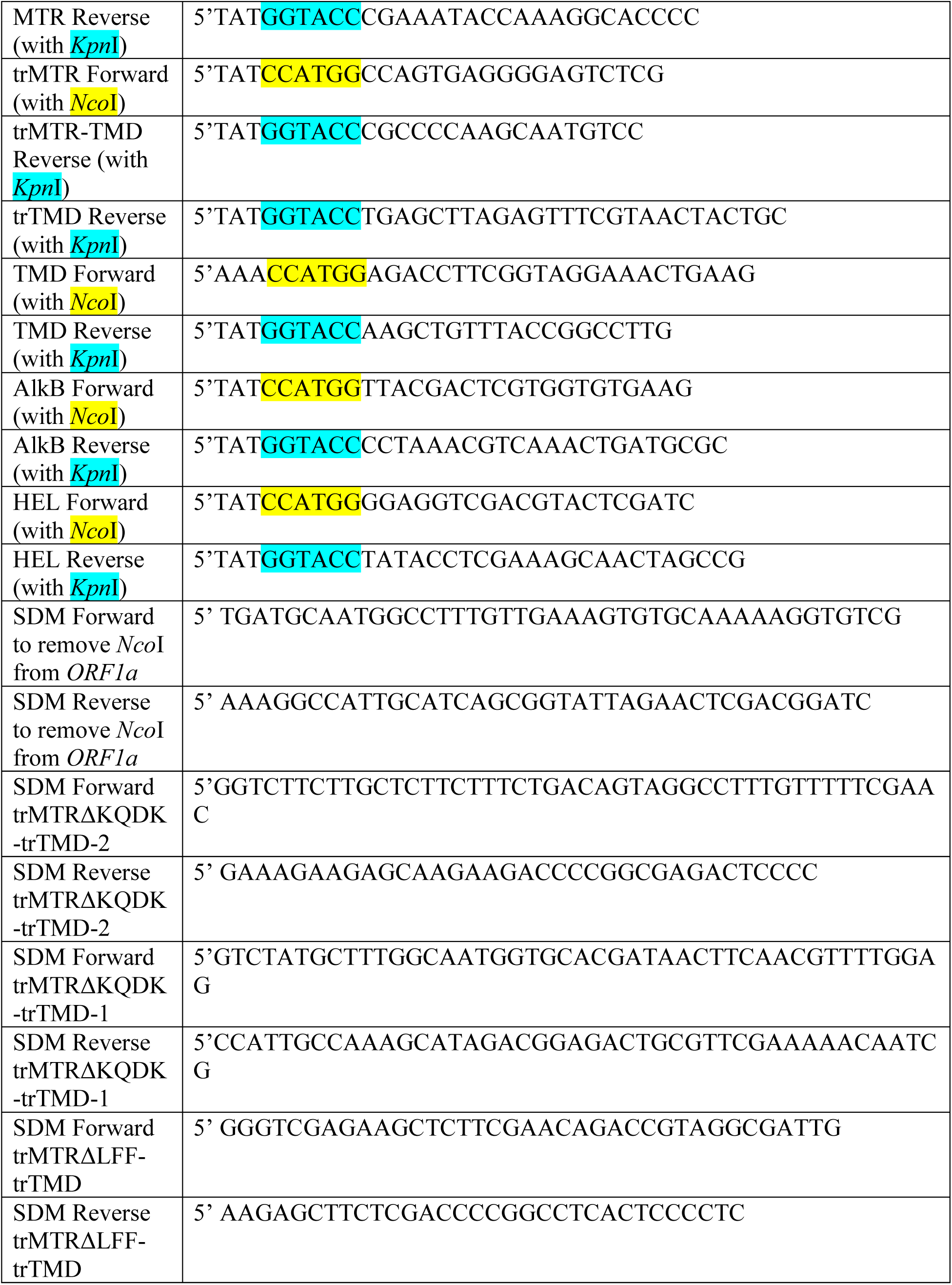
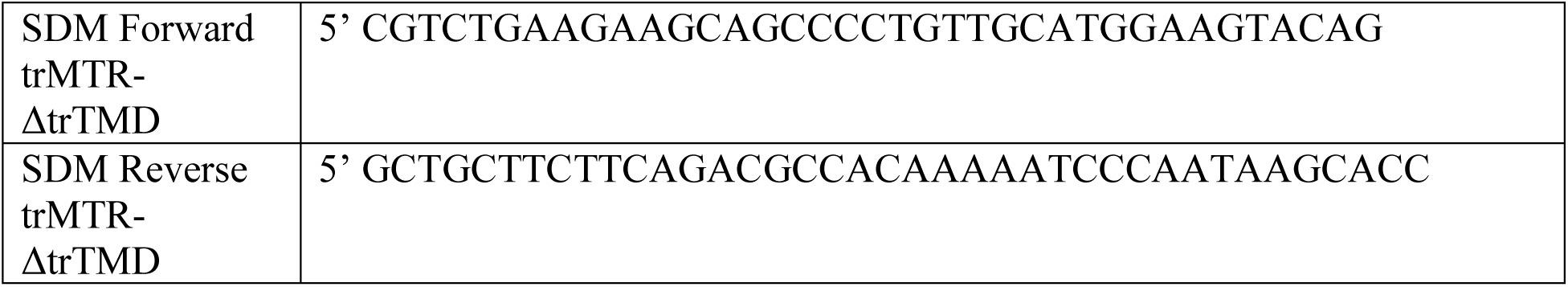
Primers utilized to clone gene expression constructs and conduct site-directed mutagenesis (SDM).

